# Caloric restriction: A potential approach for mitigating neuronal damage: Lesson from cellular model of Alzheimer disease

**DOI:** 10.1101/2024.12.11.628076

**Authors:** Apoorv Sharma, Monika Bhardwaj, Vijay Kumar, Asimul Islam, Hridayesh Prakash

## Abstract

Alzheimer’s disease is a neurodegenerative disorder and characterized by amyloid beta accumulation, synaptic dysfunction, and oxidative stress, lacks effective therapies. Caloric restriction mimetics such as fisetin and chlorogenic acid, natural polyphenols with antioxidant and autophagy-inducing properties, show promise in mitigating age-related diseases. This study investigates their neuroprotective effects against amyloid beta induced toxicity in differentiated human neuroblastoma SHSY5Y cells. Amyloid beta exposure disrupted redox homeostasis, impaired autophagy, induced mitochondrial dysfunction, and exacerbating neuronal degeneration. Fisetin and chlorogenic acid treatments reversed these deleterious effects by restoring redox balance, suppressing reactive oxygen species and upregulating critical antioxidant enzymes like SOD1, GSR, and catalase. These compounds also attenuated amyloid beta induced mitophagy via reduced PINK1 expression and restored mitochondrial fusion by upregulating Mfn2. Autophagy-related pathways were significantly modulated, evidenced by increased AMPK and decreased mTOR mRNA levels, alongside elevated expression of ATG101, ATG13, ULK1, P62 and reduced ATG5 levels. Docking studies also revealed binding of fisetin and CGA within the binding pockets of AMPK and FKBP12 supporting their interaction. Furthermore, fisetin and CGA improved synaptic integrity by upregulating PSD95 and synaptophysin and reducing acetylcholinesterase expression. These findings highlight their potential in ameliorating amyloid beta induced neuronal toxicity through autophagy activation, synaptic preservation, and mitochondrial function enhancement. While this study demonstrates the transcriptional impact and binding affinities of these caloric restriction mimetics further translational and biophysical analyses are required to elucidate their mechanisms and confirm their therapeutic viability. This research underscores the potential of fisetin and CGA as neuroprotective agents, offering promising therapeutic avenues for combating related Alzheimer’s disease neuropathies.

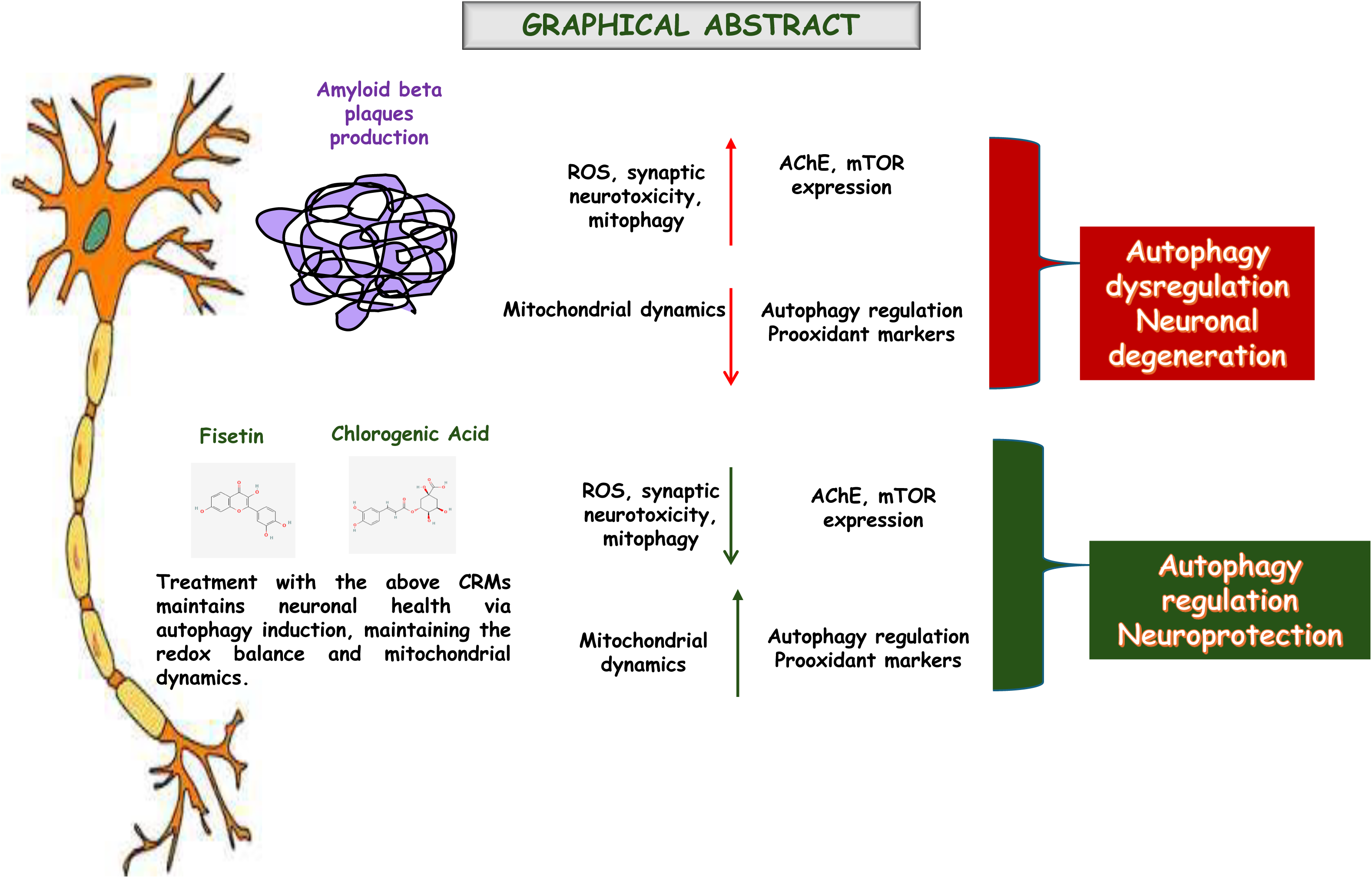

## Introduction

Autophagy, a conserved catabolic process that helps the body replenish nutrients under starving conditions, is crucial in maintaining neuronal health. It aids in the removal of toxic proteins and damaged organelles, which are the primary cause of age-related neurodegenerative disorders like Alzheimer’s, Parkinson’s, Huntington’s, etc [1]. In the case of Alzheimer’s Disease (AD), the accumulation of Amyloid beta plaques leads to severe neuronal degeneration and synapse loss. As the person ages, the process of autophagy starts declining, and the toxic clumps of misfolded amyloid beta plaques keep accumulating between neurons, exacerbating the disruption of neural networks and neuronal loss [2]. The neuronal loss in AD is irreversible. Autophagy, with its role in regulating neuronal survival [3], is not just a process but a key to understanding and potentially enhancing the body’s defense against neurodegenerative disorders. At autophagy initiation, the Unc51-like kinase 1 (ULK1) forms a complex with essential autophagy proteins – ATG101, ATG13 and FIP200/RB1CC1. This complex is necessary for autophagosome formation, regulated by autophagy regulatory proteins AMPK & mTORC [4]. The production of autophagic vesicles depends on another autophagy gene, ATG5. Autophagy can be completely inhibited or downregulated when ATG5 is knocked down, indicating that ATG5 is essential for autophagy [5]. p62, a scaffold protein, promotes autophagy degradation by binding to LC3. p62 serves as an autophagy receptor in clearing unwanted protein molecules and aggregates [6].

When core autophagy-related genes such as ATG101, ATG13, ULK1, p62, and ATG5 are dysregulated, it can result in faulty autophagosome formation, impaired lysosomal degradation, and disrupted mitophagy [7]. This, in turn, hinders cellular clearance mechanisms, accumulating toxic protein aggregates and organelle dysfunction, ultimately culminating in neuronal death. The severity increases as more neurons die, and the collapse of the synaptic network leads to cognitive and motor decline. In the early AD stage, the neurons most affected are primarily located in the entorhinal cortex and the hippocampus. In the later stages of AD, the neurodegeneration spreads extensively throughout the brain, including the areas responsible for vital body functions. The brain experiences atrophy, a significant loss of brain mass. Neurons in the entorhinal cortex and hippocampus are highly connected and rely on extensive synaptic communication. These regions are metabolically active and particularly vulnerable to the disruptions caused by amyloid-beta plaques [8]. The excitatory neurons, which are glutamatergic, cholinergic, and pyramidal neurons, are significantly impacted by amyloid-beta plaques. Amyloid-beta interferes with glutamate uptake and disrupts acetylcholine release, leading to excitotoxicity (overactivation of receptors such as NMDA) and impairments in cognitive functions like memory and attention, respectively, eventually resulting in neuronal death [9]. Loss of cholinergic signaling is a significant factor behind the cognitive decline seen in AD [10].

In AD, excessive mitochondrial fission results in neuronal death and compromised mitochondrial function [11]. A crucial mitochondrial function, mitochondrial biogenesis, that helps the cells to increase their mitochondrial mass via the fusion/fission process is impaired in AD [12]. While treatments aim to slow progression, they cannot stop or reverse the neurodegenerative damage. Therefore, finding new treatments and prevention strategies are required to significantly improve the quality of life for individuals with neurodegenerative diseases.

Calorie restriction (CR) is a dietary intervention proven to extend lifespan and improve health span in various animal models, including rodents, primates, and lower organisms [13]. Caloric restriction mimetics (CRMs) are polyphenol compounds that mimic the positive effects of caloric restriction (CR) without reducing caloric intake. Essential criteria of CRMs are autophagy induction, extending lifespan, reducing the severity of age-related illnesses and lowering protein acetylation without necessitating actual caloric restriction [14]. In this study, we have focused on exploring the potential of CRMs - Fisetin and chlorogenic acid to enhance autophagy. Both fisetin and chlorogenic acid have shown a potential to enhance autophagy, a particularly intriguing finding that warrants further investigation. [15–17], immune responses, improving metabolic functions, and antioxidant activities [18–20], but their specific effects on autophagy in AD are unclear. Both compounds influence autophagy indirectly through various signaling pathways, but their role as autophagy inducers against amyloid beta accumulation has not been explored yet.

Autophagy dysfunction is a critical factor in Alzheimer’s disease, contributing to the accumulation of amyloid-beta plaques and tau tangles, which drive neurodegeneration [2]. CRMs’ potential to enhance autophagy is a promising development, offering a hopeful outlook for the future of AD treatment. The findings of this study could lead to the development of new treatment strategies that focus on enhancing autophagy, and AMPK activators and mTOR inhibitors are promising candidates for AD treatment.

## Materials and methods

### Chemicals

The human Aβ1-42 peptide was purchased from Sigma-Aldrich Chemical Co. USA. All trans-retinoic acid (tRA), Dulbecco’s Modified Eagle’s medium, fetal bovine serum (FBS), and 100X-antibiotic/antimycotic solution, 3-(4,5-dimethylthiazol-2-yl)-2-5-diphenyltetrazolium bromide (MTT), were purchased from Gibco USA. RNeasy Mini kit for RNA extraction from Qiagen. iScript cDNA synthesis kit and SsoAdvanced^TM^ Universal SYBER Green supermix were bought from Biorad. transRetinoic-acid bought from Sigma-Aldrich, Chemical Co. USA. Fisetin and chlorogenic acid were purchased from TCI Chemicals (India) Pvt. Ltd. All the experiments were performed using autoclaved Milli-Q water. The primer sequence was synthesized and bought from Barcode Biosciences, India. The plasticware for cell culture was purchased from Nest Biotechnology Co Ltd.

### Aβ Aggregation

The powdered Aβ1-42 peptide was mixed in a stock solution at a concentration of 1 mg/ml in sterile 0.9 % normal saline, then put to incubation at 37 °C for 4 days to cause aggregation [21] and stored at-80°C.

### Cell Cultures

Human Neuroblastoma cells (SHSY5Y) were purchased from the National Centre for Cell Sciences, Pune, India. The cells were grown in a DMEM high glucose (4.5 g/L) medium with 12% FBS, 0.2% sodium bicarbonate, and 1% antibiotic/antimycotic solution. The cells were kept at 37°C in a humidified 5% CO_2_ environment. The medium was changed twice per week. SH-SY5Y cells were differentiated with All trans-retinoic acid (10^-5^ M) in 2% complete DMEM for five days to prevent unnecessary cell proliferation and develop a mature neuronal phenotype. The differentiated medium was changed twice. After differentiation, the cells were then subjected to experimental treatments. Preliminary experiments with increasing Aβ 1-42 concentrations were conducted to choose the half-maximal inhibitory concentration (IC_50_).

### Metabolic activity

The metabolic activity / viability of cells was based on their ability of viable cells to transform soluble MTT {3-(4,5n-dimethylthiazol-2-yl)-2-5-diphenyltetrazolium bromide} into an insoluble purple colored formazan crystal. Cells were seeded in 96-well plates (5000 cells/well). The cells were proliferated and differentiated as described above. After that, cells were treated with different concentrations of Chlorogenic acid (25μM-200μM), Fisetin (15μM – 120μM), and Aβ1-42 (1μM - 2μM) for 24 h and 48 h. After the treatments, the metabolic activity of the cells of different groups was analyzed by incubating them for 4h at 37°C with MTT (0.5mg/mL in DMEM) in a 5% CO_2_ incubator. After 4 h, the MTT solution was removed, and formazans were dissolved in DMSO (100μL per well). The formazans were measured at an absorbance of 570nm on BeneSphera ELISA microplate reader E21, India. The percentage of cell viability was calculated in comparison with the untreated group.

### Annexin V – FITC Apoptosis Detection Kit

The apoptosis-induced morphological changes in differentiated SHSY5Y cells after treatments were detected using Annexin V-FITC and PI apoptosis detection kit (eBiosciences^TM^ Invitrogen ThermoFisher SCIENTIFIC). Annexin V is a calcium-dependent phospholipid-binding protein and is an early apoptosis marker due to its binding with phosphatidylserine, while propidium iodide (PI) is considered a late-stage apoptosis marker due to it entering the cells once cell membrane integrity is compromised. FITC labelling allows simple direct detection by FACS analysis. Counterstaining by PI enables the discrimination of apoptotic cells. The cells were seeded onto a 24-well culture plate and were differentiated as described above. On the 5^th^ day of differentiation, the cells were treated with Aβ1-42, Fisetin & chlorogenic acid for the next 24 h (both alone and co-treated). After 24 h of incubation, the cells were trypsinized and subjected to vital double staining with Annexin V-FITC and PI according to the manufacturer’s protocol. The different experimental groups were then analysed for morphological changes in the flow cytometry analyser.

### Intracellular ROS Measurement

Intracellular ROS generation induced by Aβ1-42, Fisetin & chlorogenic acid in differentiated SHSY5Y cells was examined using a fluorescence dye Dichloro-dihydro-fluorescein diacetate (DCFH-DA) by flow cytometry. The cells were seeded onto a 24-well culture plate and were differentiated as described above. On the 5^th^ day of differentiation, the cells were treated with Aβ1-42, Fisetin & chlorogenic acid for the next 24 h (both alone and co-treated). After 24 h of incubation, the cells were washed with DPBS and incubated with DCFDA dye in serum-free DMEM (5μM working concentration) for 20 minutes in the dark at 37C. After that, the cells were again washed with DPBS and trypsinized. The ROS generation was measured at excitation and emission wavelengths of 488 nm and 535 nm, respectively.

### Real-time quantitative Reverse Transcription Polymerase Chain Reaction (RT-**qPCR)**

Total RNA was extracted according to the manufacturer’s protocol of the RNeasy Mini Kit (Qiagen-74104). RNA was quantified using Nanodrop. cDNA was prepared using the iScript cDNA synthesis kit (Bio-Rad, 170-8890). Real-time qPCR was performed using SsoAdvanced^TM^Universal SYBER Green supermix (Bio-Rad, 1725270). The RT-qPCR protocol was as follows: the first holding stage at 95 °C for 3min; followed by polymerase activation and DNA Denaturation stage at 95 °C for 15s, Annealing/Extension at 60°C for 20s, (40 cycles total). The measured data were normalized to the β-Actin level using the 2^-ΔΔCT^ method and expressed as the fold change with respect to the mean of the control group. Primer sequence is given in table 1.

### Protein-Ligand Docking

Using the UCSF Chimera, the mirror chains and other components of the receptor protein (3FAP and 4CFE) were eliminated. Protein preparation entails adding hydrogen atoms, charges, energy minimization, and the Dunbrack 2010 rotamer library to replace the incomplete side chains. The standard residues in protein preparation are charged using AMBER ff14SB, while the other residues are charged using Gasteiger. The charges are calculated using the ANTECHAMBER algorithm in both approaches. (https://doi.org/10.1080/07391102.2024.2431189)

The protein-ligand docking calculation was performed using the Autodock Vina program. The selected ligands (fisetin and CGA) were downloaded from the PubChem database, and energy was minimized using the Open Babel chemical toolbox through the MMFF94 force field. The potential atomic grid construction of spacing 0.375 Å was created, and the docking grid was adjusted around the pre-attached inhibitor of both receptor proteins. The best-docked pose based on Vina’s score was visualized using PDBsum database (https://www.ebi.ac.uk/thornton-srv/databases/pdbsum/Generate.html) and the academic version of Maestro software. (https://doi.org/10.1080/07391102.2021.2009032)

### Statistical Analysis

All the experimental data is represented as ± standard deviation (SD) of three independent experiments. The difference among multiple groups was assessed by one-way / two-way analysis of variance (ANOVA). The data was analyzed using Graph Pad Prism 9.5.0 and p < 0.05 was considered to be statistically significant.

## Result

### CRMs (Chlorogenic acid and Fisetin) mitigate beta amyloid induced metabolic arrest on differentiated SHSY5Y cells

To study the neuroprotection of CRMs, we first evaluated the cytotoxic effect of Aβ1-42 at different concentrations – 1uM, 1.25uM, 1.5uM, and 2uM- for 24 h and 48 h. We found that at 1.5uM concentration for 24 and 48 h, approximately 50% of SHSY5Y cells were metabolically arrested (Figure 1a). So, we selected 1.5uM of Aβ1-42 concentration for further experiments. The d-SHSY5Y were treated with various concentrations of chlorogenic acid (25uM, 50uM, 100uM, 200uM) and fisetin (15uM, 30uM, 60uM, 120uM) for both 24 and 48 h. Interestingly, the concentration of chlorogenic acid at 100uM and fisetin at 30uM showed a significant increase in metabolic activity (Figures 1b and 1c). Conversely, chlorogenic acid at 200uM and fisetin at 120uM contribute to the cytotoxicity in these cells. To determine the neuroprotective effect of chlorogenic acid and fisetin at 100uM and 30uM concentrations, respectively, against Aβ1-42, we did a co-treatment of chlorogenic acid (100uM) plus Aβ1-42 (1.5uM) and fisetin (30uM) plus Aβ1-42 (1.5uM) for 24 h shown in Figure 1d and 1e, respectively. We found that cotreatment with chlorogenic acid and fisetin significantly reduces the cytotoxic effect of Aβ1-42 and protects the cells from undergoing metabolic arrest.

**Figure 1:**
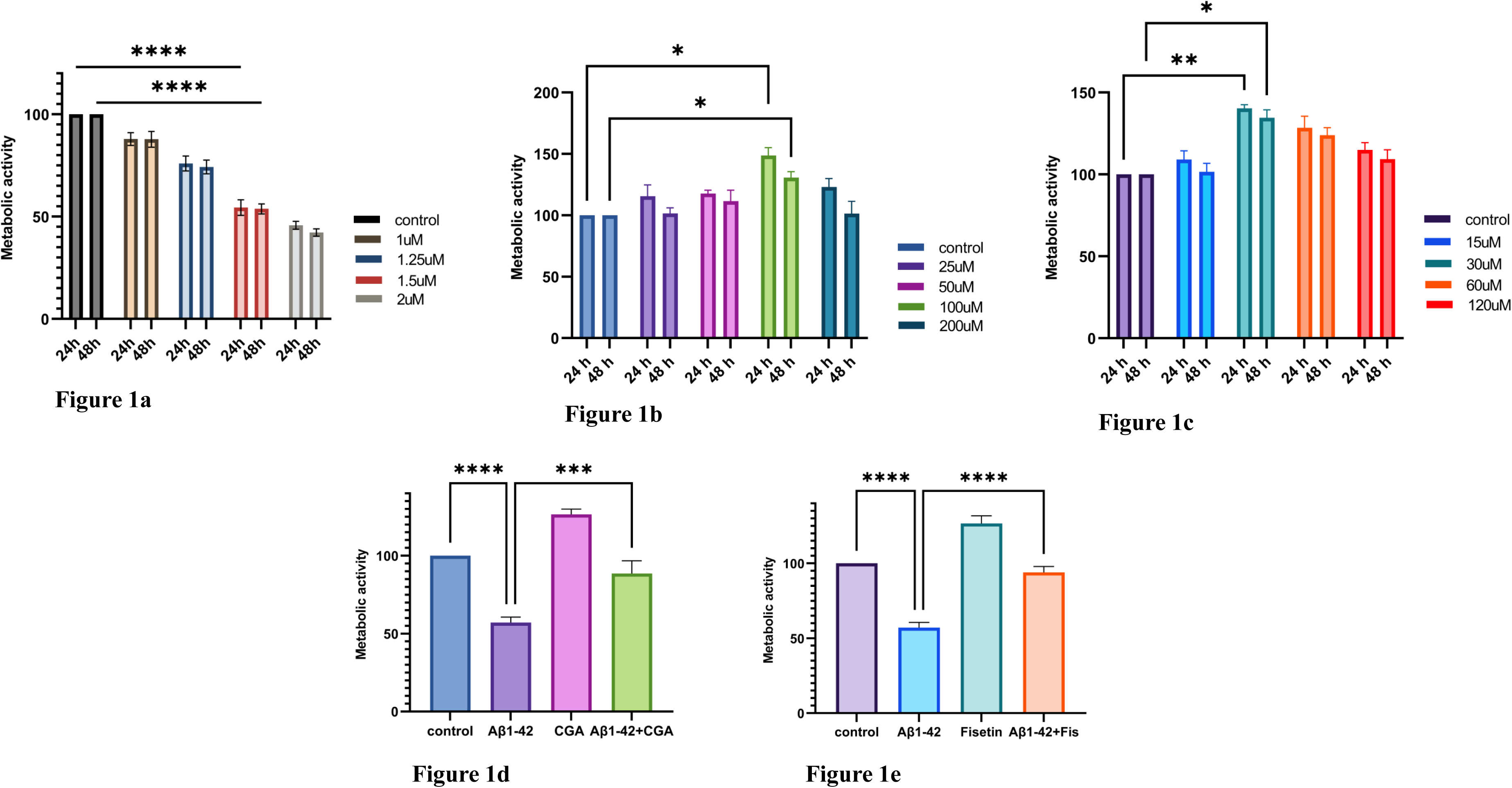
Metabolic activity: Figures 1a, 1b, and 1c show the metabolic activity of d-SHSY5Y after exposing the cells to amyloid beta (Aβ1-42), Chlorogenic Acid (CGA), & Fisetin (Fis), respectively, at different concentrations for 24 h and 48 h. In Figure 1a, the amyloid beta treatment significantly stopped the metabolic activity of d-SHSY5Y cells and caused approximately 50% metabolic arrest at 1.5µM concentration. In Figure 1b & 1c, the CGA and Fisetin significantly increased the metabolic activity at 100µM and 30µM, respectively. Figure 1d and 1e show the neuroprotective effect of CGA at 100µM and Fisetin at 30µM, respectively by showing a significant increase in metabolic activity of differentiated SHSY5Y cells in presence of Aβ1-42 at 1.5µM for 24 h. All tests were done in triplicates The data was analyzed using Graph-pad prism 9.5.0. ANNOVA two-way was applied with +/- SD, n=3, (∗ for P<0.05, ** for P< 0.01, ***for P< 0.001, **** for P<0.0001).

### Chlorogenic acid and fisetin induces autophagy in Aβ1-42 treated d-SHSY5Y Cells and rescue the neuronal cells from synaptic toxicity

To access the autophagy induction ability of chlorogenic acid and fisetin in the presence of Aβ1-42, we evaluated the fold change in mRNA expression of autophagy regulatory proteins (AMPK & mTORC), autophagy genes of ULK complex (ATG13, ATG101, ULK1), ATG5 and p62. We found that Aβ1-42 significantly upregulated mTOR mRNA levels and decreased the expression levels of autophagy genes, which are AMPK, ATG101, ATG13, and ULK1, as shown in Figure 2. Co-treatment of chlorogenic acid with Aβ1-42 and fisetin with Aβ1-42 significantly reduced the expression of mTORC to nearby control values, as shown in Figure 2a. This finding is particularly critical given that mTOR functions as a central regulatory node in cellular metabolism and autophagy, effectively inhibiting these processes when activated [22]. In contrast, AMPK acts as an energy sensor, promoting autophagy under stress conditions by inhibiting mTOR signaling, thereby fostering cellular homeostasis [23]. Notably, while Aβ1-42 inhibited the expression of autophagy-related genes and AMPK—the key signal that drives autophagy— treatment with chlorogenic acid and fisetin not only countered this suppression but also enhanced the mRNA expression levels of AMPK and critical autophagy genes such as ATG101, ATG13, and ULK1, as shown in figure 2b, 2c, 2d, and 2e, respectively. However, on checking the expression of other autophagy maker genes-p62 and ATG5, Aβ1-42 decreased the expression of p62 and increased the expression of ATG5 as shown in figure 2f and 2g, respectively.

**Figure 2:**
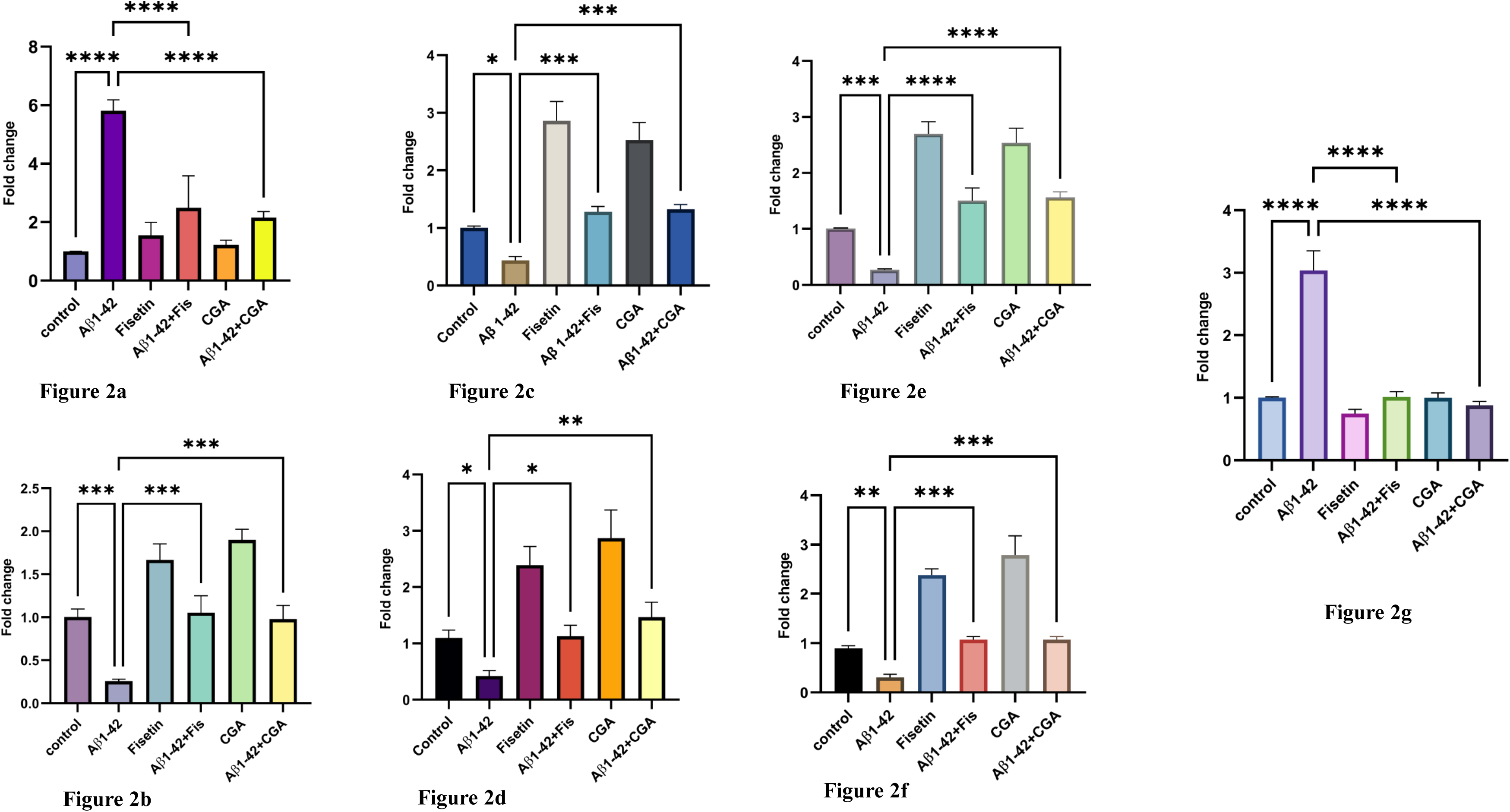
Fold Change of mRNA expression of autophagy marker genes: Figures 2a, 2b, 2c, 2d, 2e, 2f, and 2g show fold change of mTORC, AMPK, ATG101 ATG13, ULK1, p62 and ATG5 respectively. The amyloid beta treatment significantly decreased the expression of AMPK, ATG101, ATG13, ULK1, and p62 when compared to control. However, the treatment with CGA and Fisetin significantly induced the expression of autophagy genes. The amyloid beta treatment significantly increased the expression of mTORC and ATG5, compared to control, which was significantly brought down to normal levels after treating the cells with Fisetin and CGA. The data was analyzed using Graph-pad prism 9.5.0. ANNOVA one-way was applied with +/- SD, n=3, (∗ for P<0.05, ** for P< 0.01, ***for P< 0.001, **** for P<0.0001).

Dysregulation of autophagy is implicated in synaptic dysfunction [24]. Despite the critical role of neuronal autophagy in regulating synaptic architecture [25], the specific mechanisms through which autophagy modulates synaptic integrity remain underexplored. Evidence suggests that autophagy maintains synaptic homeostasis by facilitating the degradation of damaged synaptic vesicles as needed [26–28]. In this study, we aimed to elucidate the effects of chlorogenic acid and fisetin on Aβ1-42-induced synaptic toxicity in differentiated SH-SY5Y cells by examining the expression of synaptic markers through quantitative reverse transcription polymerase chain reaction (RT-qPCR). Our RT-qPCR results revealed that treatment with chlorogenic acid and fisetin (CRMs) significantly increased the mRNA transcripts of PSD95 and Synaptophysin (SYP) compared to the control group as shown in Figure 3a and 3b, respectively. In contrast, Aβ1-42 treatment led to a marked reduction in the mRNA levels of these synaptic markers relative to the control group. Notably, co-treatment with CRMs and Aβ1-42 significantly mitigated synaptic toxicity, restoring mRNA transcript levels of both PSD95 and SYP compared to the Aβ1-42-only treated group, as shown in Figure 3a and 3b, respectively.

**Figure 3:**
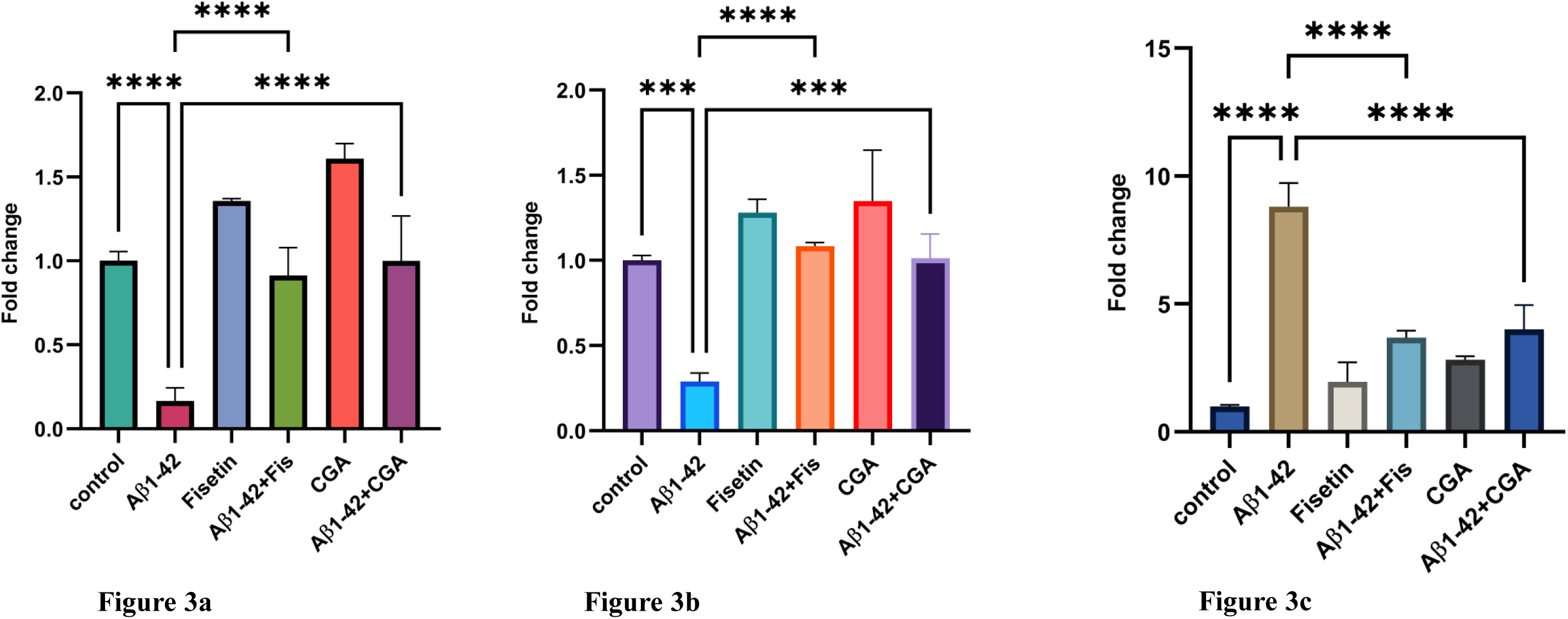
Fold Change of mRNA expression of synaptic markers and AD neuropathy marker: Figures 3a, 3b, and 3c represent the mRNA fold change of DLG4(PSD95), SYP, and AChE. The gene expression of synaptic markers was significantly reduced under amyloid beta treatment; however, on Fisetin and CGA treatment, the expression levels were brought to normal. The AD neuropathy marker AChE was significantly induced under amyloid beta toxicity. However, the Fisetin and CGA significantly ameliorated the expression of AChE and reduced the toxicity of amyloid beta. The data was analyzed using Graph-pad prism 9.5.0. ANNOVA one-way was applied with +/- SD. n=3,(***for P< 0.001, ****for P<0.0001).

Further, we evaluated the mRNA expression levels of acetylcholinesterase (AChE) across the different experimental conditions. Our findings demonstrated that Aβ1-42 treatment elicited an elevated expression of AChE, indicating a substantial increase compared to the control group, as shown in Figure 3c. Elevated AChE levels are recognized as a hallmark of Alzheimer’s disease (AD). Importantly, co-treatment with CRMs and Aβ1-42 resulted in a significant decrease in AChE expression compared to the Aβ1-42 group alone, suggesting that these CRMs may confer protective effects against AD pathology. In addition, autophagy induction necessitates the clearance of damaged mitochondria, a process known as mitophagy [29]. Thus, we assessed the expression levels of Mfn2 and PINK1 to elucidate the underlying mechanisms further.

### Chlorogenic acid and Fisetin Treatment encounter Aβ1-42-Induced Oxidative Stress and mitochondrial dysfunction in differentiated SHSH5Y

To assess the redox imbalance induced by Aβ1-42, we measured intracellular reactive oxygen species (ROS) levels (Figure 4a). Our findings indicate that Aβ1-42 disrupts the redox homeostasis of SHSY5Y cells, leading to increased intracellular ROS accumulation when compared to the control group (Figure 4a). Conversely, the treatment with chlorogenic acid and fisetin mitigated the harmful effects of Aβ1-42, maintaining ROS levels similar to those of the control group. Interestingly, when SHSY5Y cells were treated with chlorogenic acid and fisetin alone, there was a marked reduction in ROS generation compared to the control (Figure 4a). The flow cytometer data of ROS generation is given in figure 4b, which demonstrated that Aβ1-42 treatment significantly induced ROS generation to 78.6% when compared to control (58.3%). However, co-treatment of fisetin significantly reduced the ROS generation to 56.1% & with CGA it decreases to 69%. To further investigate the neuroprotective effects of these compounds against Aβ1-42-induced oxidative stress, we analyzed the fold changes of several antioxidant genes, including catalase, glutathione reductase (GSR), heme-oxygenase 1 (HO-1) and superoxide dismutase (SOD1), shown in Figure 4c, 4d, 4e and 4f, respectively. Our results demonstrated that fisetin and chlorogenic acid significantly enhanced the expression of antioxidant genes relative to the control and mitigated the detrimental effects of Aβ1-42 by sustaining the expression of these antioxidant genes and helping to maintain redox homeostasis in neuronal cells.

**Figure 4:**
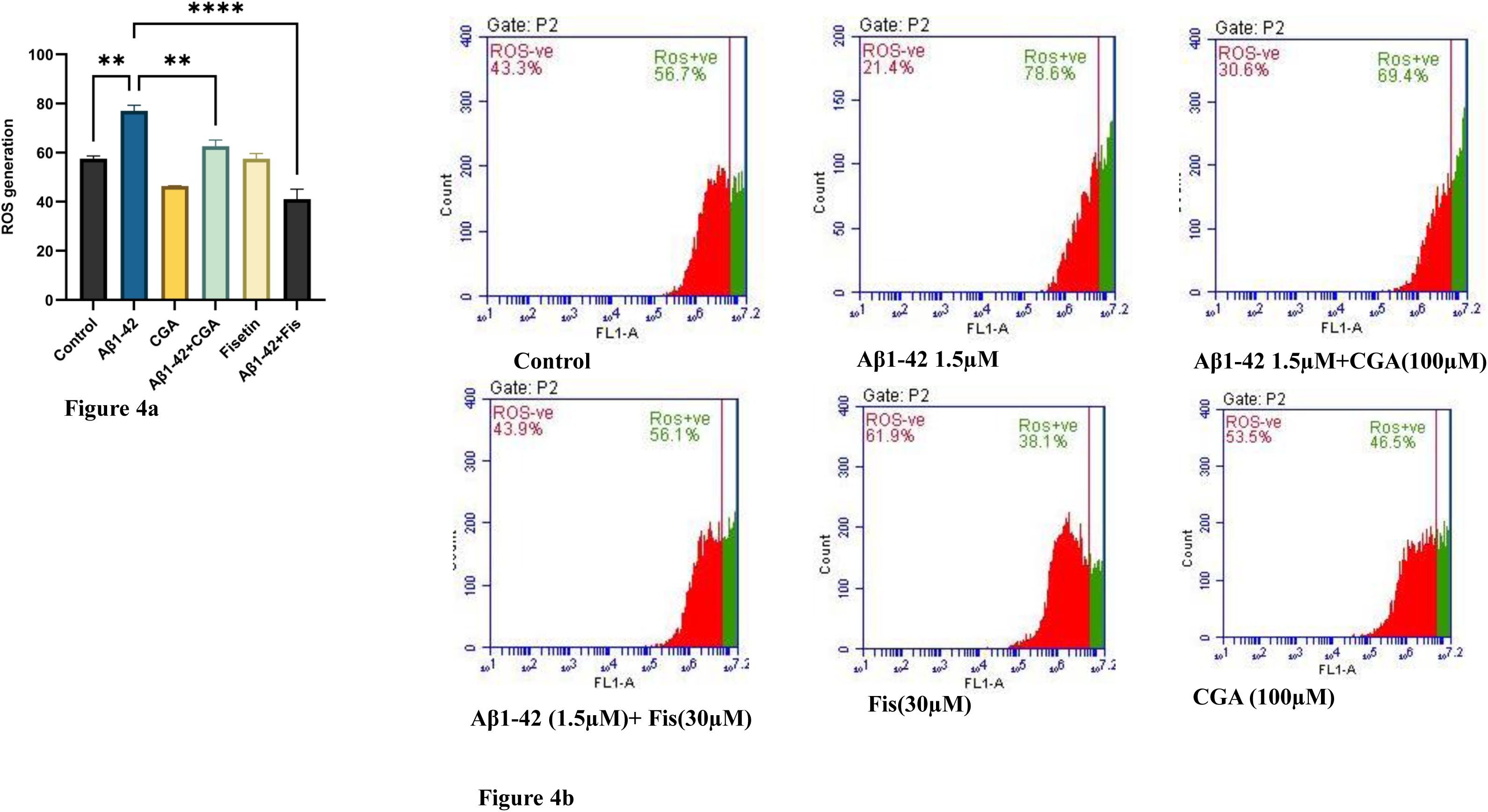
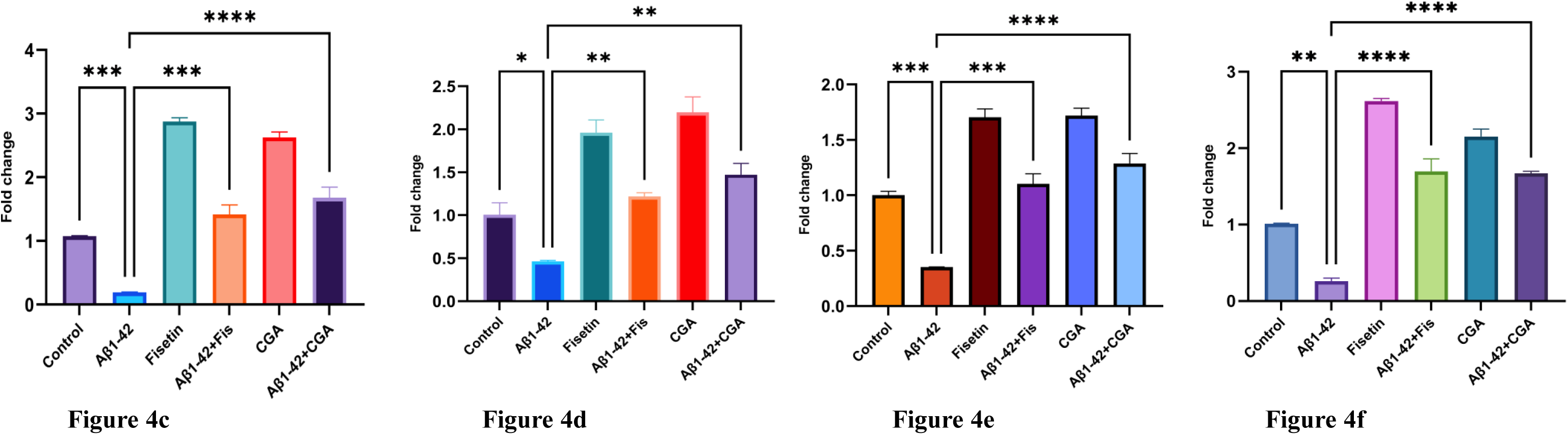
Shows ROS generation and Fold Change of mRNA expression of prooxidant markers in d-SHSY5Y cells: Figures 4a, 4b, represents the ROS generation by flow cytometer data. Figure 4c, 4d, 4e and 4f show the fold change of catalase, GSR, HO-1 & SOD-1, respectively. The amyloid beta treatment significantly decreased the expression of pro-oxidant markers when compared to control. However, the treatment with CGA and Fisetin significantly ameliorated the toxic effects of amyloid beta. n=3, (∗ for P<0.05, ** for P< 0.01, ***for P< 0.001, **** for P<0.0001).

Oxidative stress plays a pivotal role in mitochondrial dysfunction, establishing a deleterious feedback loop that may culminate in neuronal death associated with AD [30]. Mitochondria, as dynamic organelles, undergo morphological and quantitative regulation mediated by fission and fusion proteins [31]. In this context, mitofusin 2 (Mfn2) facilitates mitochondrial fusion, while PINK1 is integral to the process of mitophagy. The Mfn proteins are essential for the merging of mitochondria, thereby preserving their functional integrity [32].

To investigate these dynamics, we assessed the fold changes in Mfn2 and PINK1 expression in the presence of Aβ1-42, in conjunction with evaluating the mitigating effects of chlorogenic acid and fisetin against Aβ1-42 exposure. Our results demonstrated that SHSY5Y cells exposed to Aβ1-42 exhibited significantly reduced mRNA transcript levels of Mfn2 (Figure 5a). Notably, treatment with chlorogenic acid and fisetin restored Mfn2 transcript levels to values comparable to the control group.

**Figure 5:**
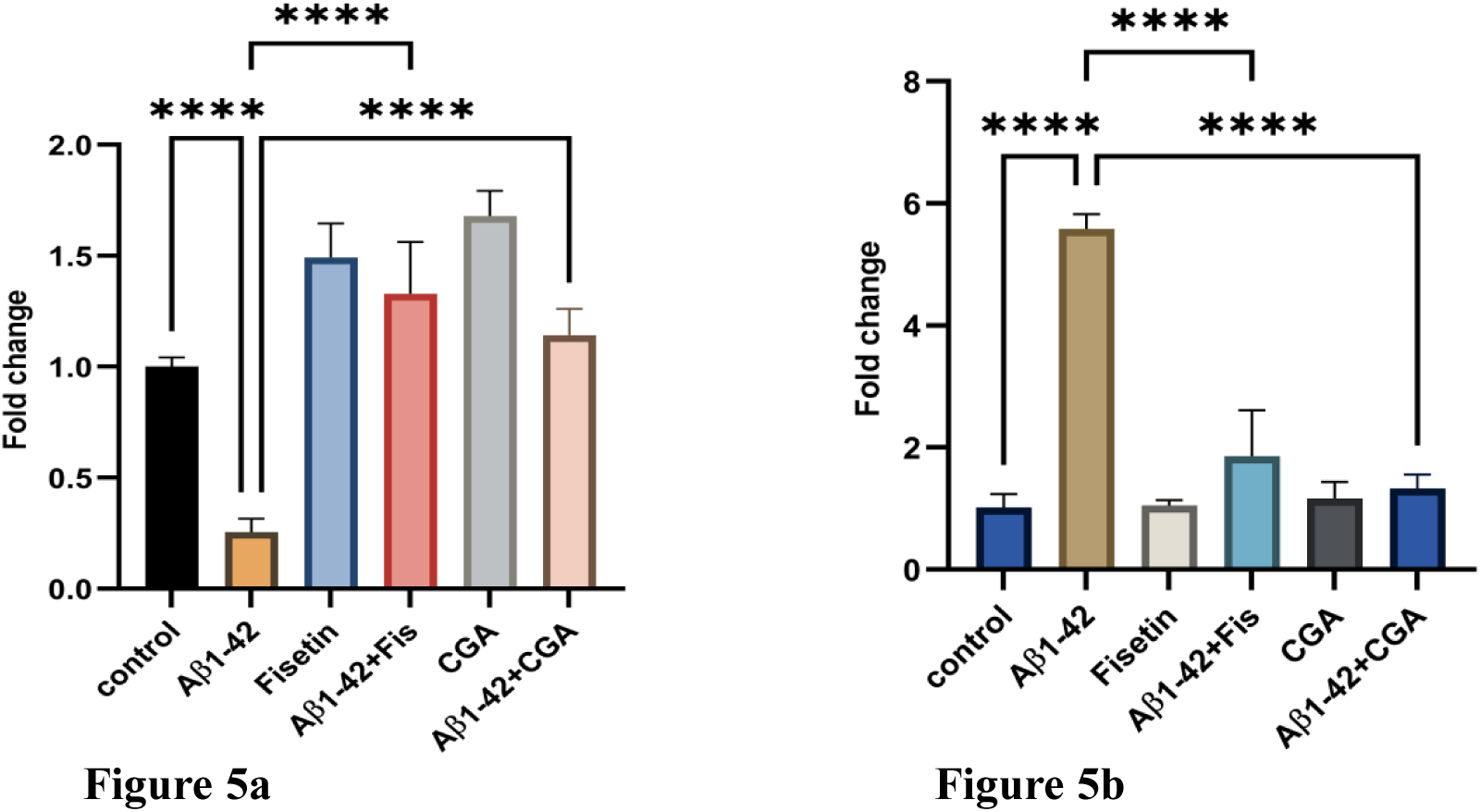
Shows the mRNA fold change of MFN2 and PINK1 in the presence of Amyloid beta, CGA, and fisetin. Figure 5a shows Amyloid beta significantly reduced the expression of MFN2, while CGA and fisetin increased the MFN2 expression significantly. In Figure 5b, Amyloid beta significantly induced the PINK1 expression, while CGA and fisetin significantly alleviated the PINK1 expression. The data was analyzed using Graph-pad prism 9.5.0. ANNOVA one-way was applied with +/- SD. n=3, (∗ for P<0.05, ** for P< 0.01, ***for P< 0.001, **** for P<0.0001).

Given the therapeutic effects of chlorogenic acid and fisetin on Aβ1-42-induced mitochondrial dysfunction, we further explored their influence on mitophagy through the evaluation of PINK1 expression fold changes (Figure 5b). The mitophagy process is critical for the elimination of dysfunctional and damaged mitochondria, processes that have been implicated in the pathogenesis of AD. Our findings revealed that Aβ1-42 significantly elevated PINK1 expression (Figure 5b) relative to the control group, suggesting a pronounced activation of mitophagy. Conversely, co-treatment of chlorogenic acid and fisetin with Aβ1-42 resulted in a significant decline in the levels of this mitophagy marker, indicating a synergistic effect in mitigating the cellular stress induced by Aβ1-42.

### Chlorogenic acid and fisetin protect confer resistance in differentiated SHSY5Y against Aβ1-42-Induced Apoptosis

Mitochondrial dysfunction is believed to play a significant role in cellular apoptosis. To assess the inhibitory effects of specific compounds against Aβ1-42-induced apoptotic cell death, flow cytometry analysis was conducted on cells stained with Annexin V-FITC and PI. Results, as shown in Figure 6 indicated, that Aβ1-42 treatment led to reduced cell viability compared to control conditions, while a marked increase in viability was observed in the groups treated with the tested compounds. Specifically, co-treatment of compounds with Aβ1-42 effectively rescued cells from apoptosis and enhanced cell viability. These findings suggest that the CRMs can effectively mitigate the deleterious effects of Aβ1-42 on cell survival.

**Figure 6:**
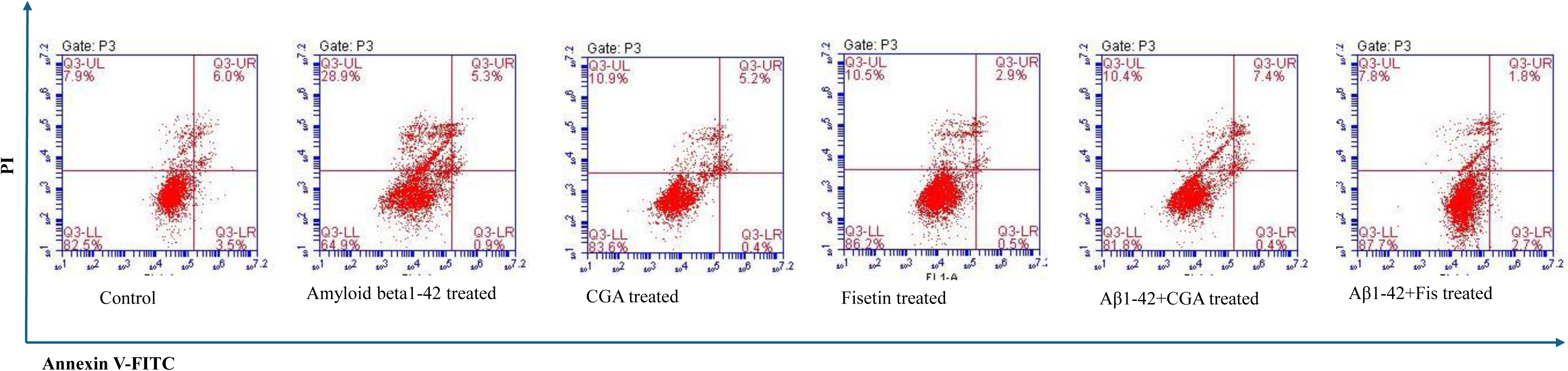
The flow cytometric analysis of apoptosis in differentiated SHSY5Ycells using AnnexinV-FITC and PI apoptosis detection kit following the treatment of Aβ1-42 (1.5μM), CGA (100μM), and fisetin (30μM) for 24 h.

### DOCKING Results

The protein-ligand interaction of fisetin and CGA was analyzed through site-specific docking on the 3FAP and 4CFE using Autodock Vina. The selected ligands were docked to the active site of both the protein receptors. According to the docking results, all the ligands fit well into the active site of FKBP12 and AMPK protein. We docked the CGA and fisetin at the FKBP12 binding pocket of mTOR [33]. In the FKBP12, the binding affinity of 3FAP-fisetin and 3FAP-CGA docked structure is −8.2 kcal mol-1 and −10.9 kcal mol-1, respectively.

In 3FAP, the CGA is bounded by hydrophobic amino acid residues (like Trp 190B, Tyr 194B, and Phe 197B), hydrogen bonds (like Tyr 26A, Asp 37A, and Thr 187B), and salt bridge (Arg 42A). On docking CGA with 3FAP, the cyclohexane ring of CGA forms three hydrogen bonds with three hydroxyl groups present at the C1, C4, and C5 with Tyr26A, Thr187B, and Asp37A, respectively. A salt bridge is formed between the carboxylic group present at C1 with a positively charged amino acid Arg42A. The caffeic acid part of CGA penetrates deep into the hydrophobic pocket of FKBP12, where it is surrounded by aromatic amino acid residues and forms hydrophobic interactions with amino acids (like Trp190B, Tyr194B, and Phe197B). The Phe197B forms a Pi-Pi stacked interaction with the aromatic ring of CGA. Glu121 forms a carbon-hydrogen bond with the hydroxyl group at C4 of the aromatic ring of CGA, as shown in Ligplot figure 7a and the 3D representation of this docked structure is shown as figure 7b.

**Figure 7:**
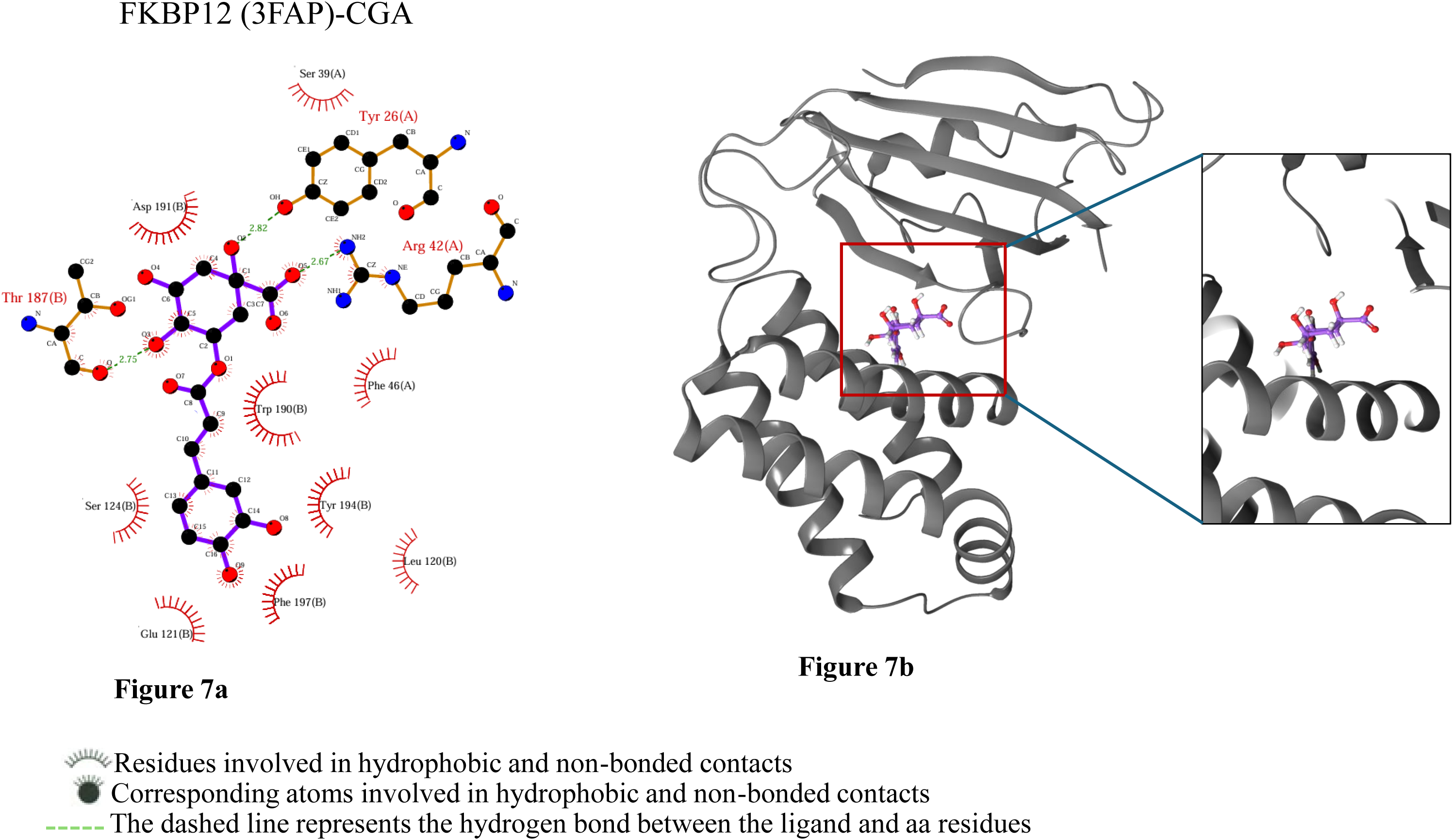
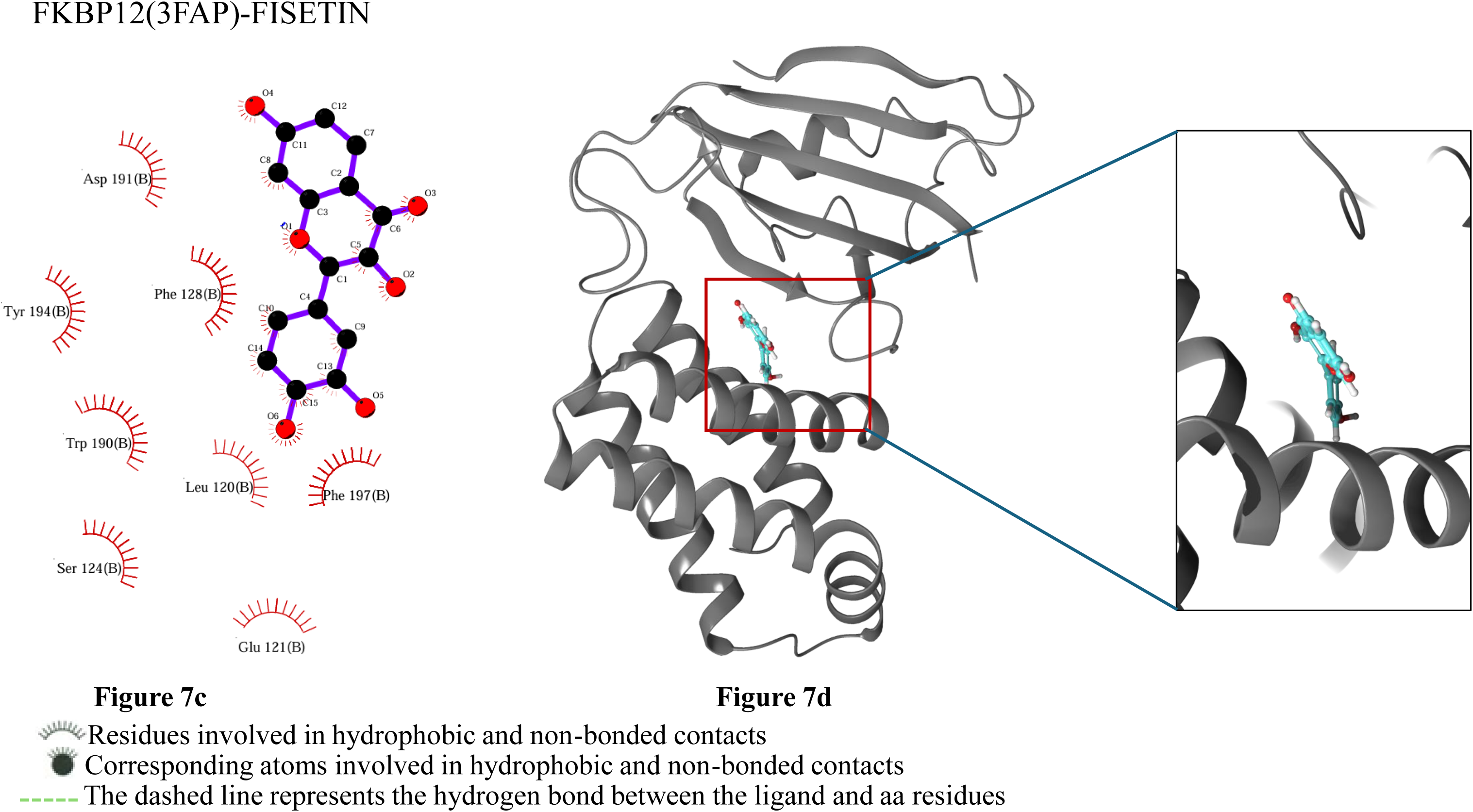
The protein-ligand interaction of CGA and fisetin with FKBP12 (3FAP). The 2D ligplot was developed from PDBsum. Figure 7a and 7b represent the docked structure of CGA with FKBP12 and fisetin with FKBP12 respectively. The 3D structure was developed from pymol, figure 7c and 7d represent the docked structure of CGA with FKBP12 and fisetin with FKBP12 respectively.

However, on docking fisetin with FKBP12, the fisetin is surrounded by the hydrophobic pocket of FKBP12, where it is surrounded by aromatic amino acid residues (like Phe 128B and Trp 190B) and engages a Pi-Pi stacking with Tyr 194B (Ligplot figure 7c). There were no hydrogen bonds, only non-bonded interactions, including hydrophobic interactions and pi-pi interactions. These interactions also reside deep within the hydrophobic pocket of FKBP12. From our docking study, we inferred that both ligands bind well within the drug-binding pocket of FKBP12, which is also a catalytic domain. The 3D interaction profile diagram of fisetin with FKBP12 is shown as figure 7d.

The structure of full-length human α2β1ү1 AMPK was downloaded from PDB: 4CFE. In this structure, the CBM is attached to the N-terminal lobe of the kinase domain and the interface between these two domains generates the binding cavity [34], where we docked the fisetin and chlorogenic acid. In the AMPK protein, the binding affinity of 4CFE-fisetin and 4CFE-CGA docked structure is −4.6 kcal mol-1 and −5.5 kcal mol-1, respectively.

CGA forms four hydrogen bonds with Gly28A, Asn48A, Ile46A, and Asp88A. All these residues belong to the kinase domain. The CGA also forms non-bonded interactions with residues Lys29A, Val24A, Phe27A, and Leu47A. All residues belong to the kinase domain, which is present at the N-terminal of the α-subunit, the C-terminal of which contains the regulatory domain. The 2D Ligplot is shown in figure 8a and the 3D representation is shown in figure 8b. Furthermore, in the 4CFE-CGA docked structure, the CGA engages hydrophobic residues like Lys 29A and Ile 46A; hydrogen bonds with Ile 46A and Asp 88A.

**Figure 8:**
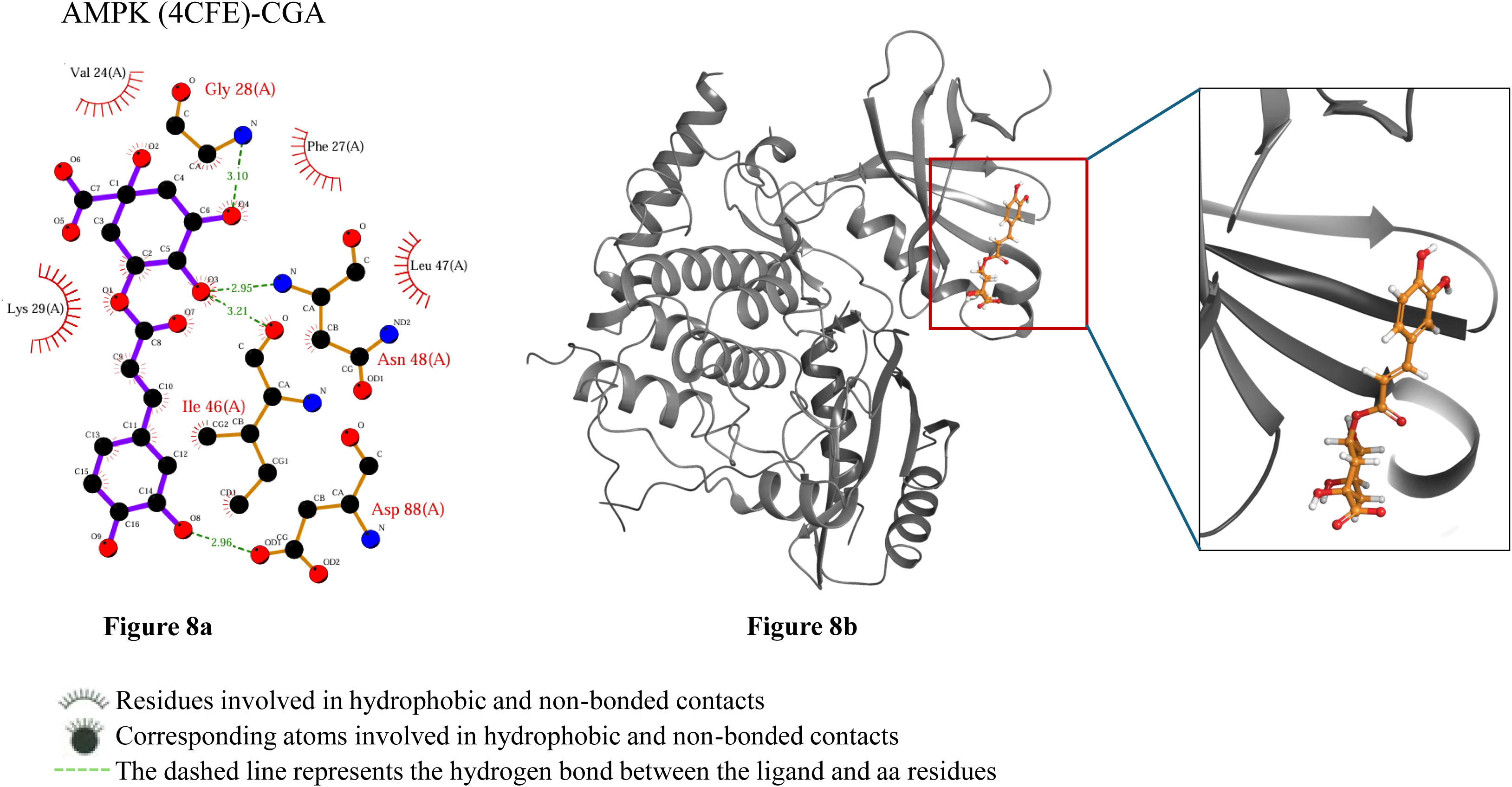
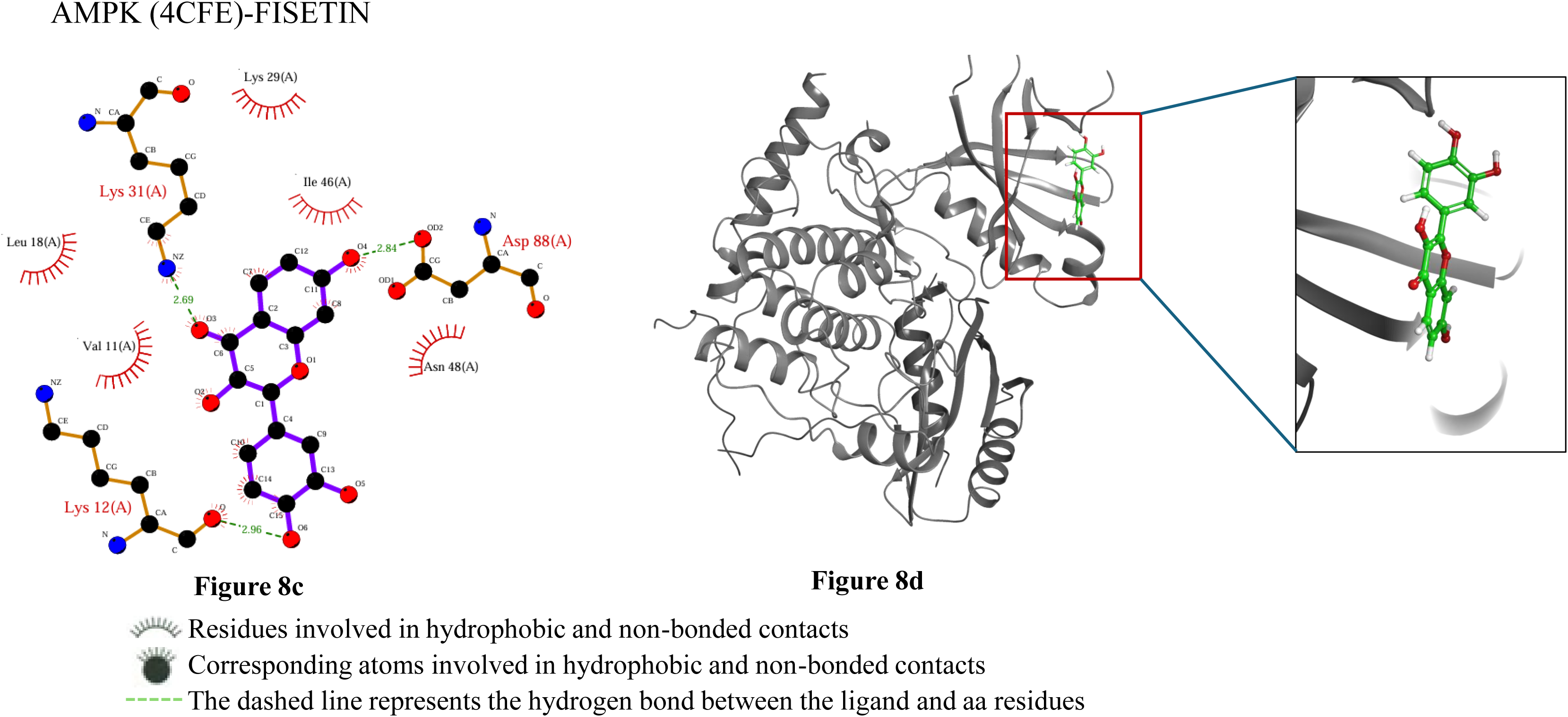
The protein-ligand interaction of CGA and fisetin with AMPK (4CFE). The 2D ligplot was developed from PDBsum. Figure 8a and 8b represent the docked structure of CGA with FKBP12 and fisetin with FKBP12 respectively. The 3D structure was developed from pymol, figure 8c and 8d represent the docked structure of CGA with FKBP12 and fisetin with FKBP12 respectively.

Fisetin, on the other hand, forms two hydrogen bonds with Asp88A and Lys12A and five non-bonded interactions with Leu18A, Val11A, Lys29A, Ile46A, and Asn48A. All these binding residues belong to the kinase domain shown in figure 8c in the form of 2D liglplot and 8d shows the 3D representation of of AMPK docked with fisetin. In the 4CFE, a few of the hydrophobic amino acid residues that circle fisetin are Val 11A, Leu 18A, and Ile 46A; the hydrogen bond interacting residues are Lys 12A, Lys 31A, and Asp 88A. Fisetin also engages in a pi-pi interaction with Phe 90A.

## DISCUSSION

The primary purpose of the present study is to explore CRM-induced neuroprotection during Aβ1-42-associated neuronal damage The CRMs were able to reverse the synaptic dysfunction and mitigate the Aβ1-42 -induced synaptic and neuronal degeneration, via autophagy induction, synaptic preservation, mitochondrial function, and maintenance of redox homeostasis.

CRMs are natural or synthetic water-soluble polyphenol compounds (12-16 phenol group) known for mimicking the beneficial effects of calorie restriction without restricting the calories. CRMs are categorized into exclusive groups of glycolytic inhibitors, sirtuin regulators, autophagy activators, lipid regulators, insulin sensitizers, antioxidants, and anti-inflammatory [35, 36]. Chlorogenic acid and fisetin are CRMs categorized as the sirtuin activators [37, 38]. They are polyphenolic compounds with free radical scavenging capabilities, making them very helpful in treating age-related illnesses. Several studies have been conducted on fisetin and chlorogenic acid, showing their ability to reduce age-related illnesses like cardiovascular, neurological, and carcinogenic problems [39–42]. However, the direct effect of these CRMs on autophagy induction in ameliorating the toxic accumulation of Aβ1-42 and rescuing the neuronal cells from synaptic toxicity and mitophagy has not been studied yet.

Our study elucidated the antioxidant, anti-apoptotic, and neuroprotective potential of activated autophagy in the context of Aβ1-42-induced toxicity in differentiated human neuroblastoma SHY5Y cells. The treatment with Aβ1-42 resulted in compromised redox status, induced mitophagy, exacerbated apoptotic cell death, and inflicted synaptic neurotoxicity in these cell lines. Aβ1-42 has been well-documented to provoke oxidative stress, neuronal dysfunction, and neuronal cell death, ultimately leading to neurotoxicity [43]. Aβ1-42 is a pivotal molecular contributor to the pathophysiology of AD [44]. Consistent with previous research, our findings demonstrated that Aβ1-42 exhibited substantial toxicity towards d-SHY5Y cells, which was reversed by chlorogenic acid and fisetin (Figure 1a-1e).

Our findings demonstrate that chlorogenic acid and fisetin induced autophagy provides neuroprotection against Aβ1-42 toxicity by increased AMPK and decreased mTOR mRNA expression. In addition, chlorogenic acid and fisetin increased the mRNA levels of ATG101, ATG13, and ULK1, essential components required for autophagosome formation. Activation of this complex occurs through AMPK-mediated phosphorylation of ULK1, which then initiates downstream autophagy processes [45]. Furthermore, our study showed that Aβ1-42 elevated ATG5 expression, which was in consistent with the previous studies, revealing that in the late stages of AD, a continual buildup of Aβ led to excessive autophagy, which resulted in neuronal dysfunction and aggravated AD symptoms [46, 47]. However, the treatment with chlorogenic acid and fisetin maintained the mRNA levels of ATG5. The reduced p62 mRNA levels under Aβ1-42 toxicity suggest autophagy dysregulation which was maintained by chlorogenic acid and fisetin treatment. These results highlight the potential of these compounds to modulate autophagy pathways and offer neuroprotection in Aβ1-42-induced cellular dysfunction.

The pathogenesis of AD correlates with neuronal dysfunction and loss of functional synapses caused by changes in synaptic proteins. PSD95 (postsynaptic density protein 95) and synaptophysin are critical proteins involved in synaptic function and neurotransmission in the brain. PSD95 is a scaffolding protein predominantly found at the postsynaptic density of excitatory synapses. It plays a crucial role in organizing and anchoring neurotransmitter receptors, such as NMDA and AMPA, at the postsynaptic membrane [48]. This organization is essential for synaptic signaling, plasticity, and neuron communication. Synaptophysin is a membrane protein located on synaptic vesicles in presynaptic terminals. It regulates neurotransmitter release by facilitating the docking and fusion of synaptic vesicles with the presynaptic membrane. Our study correlates with the previous studies that Aβ disrupts the normal function of both PSD95 and synaptophysin, leading to impaired synaptic communication [46, 49]. In our study, fisetin and chlorogenic acid attenuated the synaptotoxic effect of Aβ1-42 by inducing the expression of PSD95 and synaptophysin (Figure 3a, 3b). One of the main hallmarks of AD pathology is elevated expression of acetylcholinesterase (AChE) [50]. Our study showed Aβ1-42 treatment significantly increased the expression of AChE. Treatment with chlorogenic acid and fisetin reduced the AChE levels and mitigated the toxic effect of Aβ1-42 (Figure 3c).

Redox homeostasis is essential for proper cellular and molecular physiology functioning. Neuronal cells are particularly vulnerable to oxidative stress-mediated cell death due to their high polyunsaturated fatty acids and relatively low antioxidant defenses [51]. A substantial body of research suggests that cells rely on antioxidant enzymes to effectively cope with oxidative stress [52, 53]. We find that chlorogenic acid and fisetin reversed Aβ1-42-mediated redox imbalance in SH-SY5Y cells by decreasing the levels of reactive oxygen species (ROS) with concomitant increased expression activity of critical antioxidant enzymes, such as superoxide dismutase-1 (SOD1), glutathione reductase (GSR), catalase, and heme oxygenase-1 (HO-1) (Figure 4a-4f).

Mitochondrial dynamics, encompassing the processes of fusion and fission, play a critical role in mitochondrial function, which are disrupted by amyloid-beta (Aβ) in AD [54]. The present study demonstrated that chlorogenic acid and fisetin restored the reduced mRNA levels of the mitochondrial profusion gene Mfn2 induced by Aβ1-42 in d-SH-SY5Y cells. Mitochondrial dysfunction is another hallmark of amyloid beta induced neurotoxicity in AD patients [55].

The observed decrease in mRNA levels of mitochondrial profusion genes and the concomitant increase in reactive oxygen species (ROS) levels lead to mitophagy, which targets damaged mitochondria for degradation. Under physiological conditions, mitophagy is essential for maintaining basal mitochondrial turnover and homeostasis. The PTEN-induced putative kinase 1 (PINK1)-Parkin-mediated pathway represents one of the most extensively characterized mechanisms of mitophagy [56, 57]. Previous research has shown that acute depolarization of the mitochondrial membrane potential (Δψm) in vitro, induced by Δψm-dissipating agents, triggers Parkin-mediated mitophagy, which facilitates the removal of dysfunctional mitochondria via the autophagy-lysosomal system. Enhanced mitophagy has been documented in the brains of AD patients, correlating with elevated PINK1 expression [58]. In this study, we provided evidence that chlorogenic acid and fisetin effectively mitigated Aβ1-42-induced mitophagy by lowering PINK1 levels (Figure 5b).

Further, the increase in oxidative stress attributable to Aβ1-42 disrupts redox homeostasis, consequently leading to mitochondrial dysfunction, associated with heightened cellular apoptosis [59]. Our results suggest that Aβ1-42 induced apoptosis in d-SHSY5Y cells is mitigated by fisetin and chlorogenic acid (Figure 6). However, further investigation is warranted to elucidate the specific apoptotic pathways activated in differentiated SH-SY5Y cells treated with Aβ1-42.

Next, our docking results demonstrate that CGA and Fisetin bind strongly within the drug binding pocket of mTOR (FKBP12) and in the kinase domain of AMPK. The FKBPs (FK506-binding proteins) were initially identified as ubiquitously expressed immunophilins that mediate the pharmacological effects of naturally occurring macrolide immunosuppressants, such as FK506 and rapamycin. These proteins function as peptidyl-prolyl cis-trans isomerases (PPiases), facilitating the conversion of cis-proline residues to a less sterically constrained trans conformation. In addition, FKBPs play roles in the regulation of intracellular calcium release, gene transcription, protein translation, and cellular trafficking. Among the FKBP family, FKBP12 is the smallest member and features a basic domain, a PPiase catalytic domain, and a drug-binding pocket. FKBP12 serves as the intracellular receptor for rapamycin, a widely used inhibitor of mTOR signaling by disrupting the interaction between mTOR and Raptor at the FKBP12-binding domain (FRB) on mTOR [33, 60]. Consequently, it is conceivable that FKBP12 typically functions to repress mTOR activity and the high binding affinity of chlorogenic acid and fisetin with FKBP12, suggest that these two compounds have inhibitory effect on mTOR.

AMPK is a heterotrimeric enzyme complex composed of a catalytic α-subunit and regulatory β- and γ-subunits. The α-subunit features an N-terminal protein kinase domain along with a C-terminal regulatory domain. The γ-subunit contains four iterations of a cystathionine β-synthase domain. It is crucial for regulating energy homeostasis in eukaryotic cells and it is activated through the phosphorylation of a threonine residue (Thr-172) located within the activation loop of its kinase domain [34]. Due to its significant role in energy regulation, AMPK represents an attractive target for drug development aimed at preventing or mitigating the effects of metabolic diseases. Our docking study inferred that fisetin and CGA bind well within the kinase domain of AMPK that goes along with the previous studies of small molecule activators A-769662 [34]. But the effect of fisetin and CGA on AMPK phosphorylation still needs to be explored.

In both of the above cases, further research needs to be done about the inhibition of mTOR and activation of AMPK using these compounds as inhibitors and activators, respectively. Since we have shown the docking and transcriptional data. Translational validation still needs to be done. The conformational changes that occur in the kinase domain of AMPK and in the catalytic domain of FKBP12 during the binding process are yet to be explored. Analysing the stability of the complexes is also yet to be explored. Only after an extended biophysical characterization can we deduce whether these compounds can be categorized as potent autophagy inducers. Our findings demonstrate that treatment with fisetin and chlorogenic acid significantly attenuates the toxic effects of Aβ1-42 by inducing autophagy, preserving the synapses, maintaining the redox homeostasis and mitophagy highlighting the potential therapeutic implications of these compounds in combating AD-related neuropathies.

## Conclusion

The study underscores that chlorogenic acid and fisetin can effectively counteract the detrimental effects of Aβ1-42 by inducing autophagy, restoring synaptic health, maintaining redox status, and mitigating mitophagy. However, further research is needed to assess the efficacy and safety of chlorogenic acid and fisetin in AD animal models. Understanding these compounds’ pharmacokinetics and optimal dosing regimens could pave the way for clinical trials. Furthermore, exploring the synergistic effects of combining these compounds with existing AD therapies could enhance their therapeutic potential. Overall, harnessing the protective effects of nutraceuticals like chlorogenic acid and fisetin may lead to innovative strategies for preventing or slowing the progression of Alzheimer’s disease.

## Statements and Declarations

### Data Availability

All the data is present in the manuscript

### CRediT authorship contribution statement

**Apoorv Sharma:** The first draft of the manuscript, diagrams, data analysis and table. **Monika Bhardwaj:** Study validation, analysis and editing of manuscript. **Dr Hridayesh Prakash:** Study design, analysis, conceptualization and validation. **Dr Vijay Kumar:** Reviewed and edited. **Asimul Islam**: Reviewed and edited. All authors read and approved the final manuscript.

## Acknowledgment

We would like to express our sincere gratitude to all those who contributed to the completion of this study. We also wish to thank the authors of the studies cited in this study, whose research forms the foundation of our work. We would like to thank Puneet Kumar for providing guidance in docking study.

